# PCaDB - a comprehensive and interactive database for transcriptomes from prostate cancer population cohorts

**DOI:** 10.1101/2021.06.29.449134

**Authors:** Ruidong Li, Jianguo Zhu, Wei-De Zhong, Zhenyu Jia

## Abstract

Prostate cancer (PCa) is a heterogeneous disease with highly variable clinical outcomes which presents enormous challenges in the clinical management. A vast amount of transcriptomics data from large PCa cohorts have been generated, providing extraordinary opportunities for the molecular characterization of the PCa disease and the development of diagnostic and prognostic signatures. The lack of an inclusive collection and harmonization of the scattered public datasets constrains the extensive use of the valuable resources. In this study, we present a user-friendly database, PCaDB, for a comprehensive and interactive analysis and visualization of gene expression profiles from 77 transcriptomics datasets with 9,068 patient samples. PCaDB also includes a single-cell RNA-sequencing (scRNAseq) dataset for normal human prostates and 30 published PCa prognostic signatures. The comprehensive data resources and advanced analytical methods equipped in PCaDB would greatly facilitate data mining to understand the heterogeneity of PCa and to develop machine learning models for accurate PCa diagnosis and prognosis to assist on clinical decision-making. PCaDB is publicly available at http://bioinfo.jialab-ucr.org/PCaDB/.

## Introduction

As one of the most powerful approaches in oncology research, transcriptome profiling has been extensively used for understanding the molecular biology of cancer, drug target identification and evaluation, biomarker discovery for cancer diagnosis and prognosis, etc. over the past decade (1). Prostate cancer (PCa) is the second most frequently diagnosed cancer in men worldwide with 1,414,259 new cases and 375,304 new deaths in 2020 (2). The vast amount of the publicly available transcriptomics data for clinical samples provides extraordinary opportunities for the study of the heterogeneity in PCa, understanding the mechanisms of therapeutic resistance, as well as identification and validation of diagnostic and prognostic signatures to improve PCa management. Existing pan-cancer databases such as GEPIA (3) and cBioPortal (4) are exceptionally useful, however, only a few PCa transcriptomics datasets have been included in those databases. It remains challenging for researchers to leverage the valuable resources in their studies without a comprehensive collection and harmonization of the scattered PCa transcriptomics datasets. Moreover, advanced programming skills and deep knowledge in bioinformatics and data science are also required for conducting a comprehensive analysis and visualization of the datasets.

To fill this void, we present to the PCa research community a user-friendly database, PCaDB (http://bioinfo.jialab-ucr.org/PCaDB/), for a comprehensive and interactive analysis and visualization of transcriptomics data from PCa cohorts (Figure 1). A comprehensive bioinformatics workflow has been developed to download, process, and harmonize the gene expression data and sample metadata for 77 PCa transcriptomics datasets with a total of 9,068 samples from public data repositories. A single-cell RNA-sequencing (scRNAseq) dataset for normal human prostates has been included in PCaDB, allowing for the investigation of gene expression in different cell types of prostate (5). Moreover, PCaDB included 30 published PCa prognostic signatures which have been analyzed in a previous study for a comprehensive evaluation of the performances of machine learning models and prognostic signatures (6). The harmonized data resources and analytical functions provided in PCaDB will significantly promote the studies for better understanding of the molecular basis of PCa and the discovery of biomarkers and molecular signatures for PCa diagnosis and prognosis to improve PCa management.

**Figure 1.**
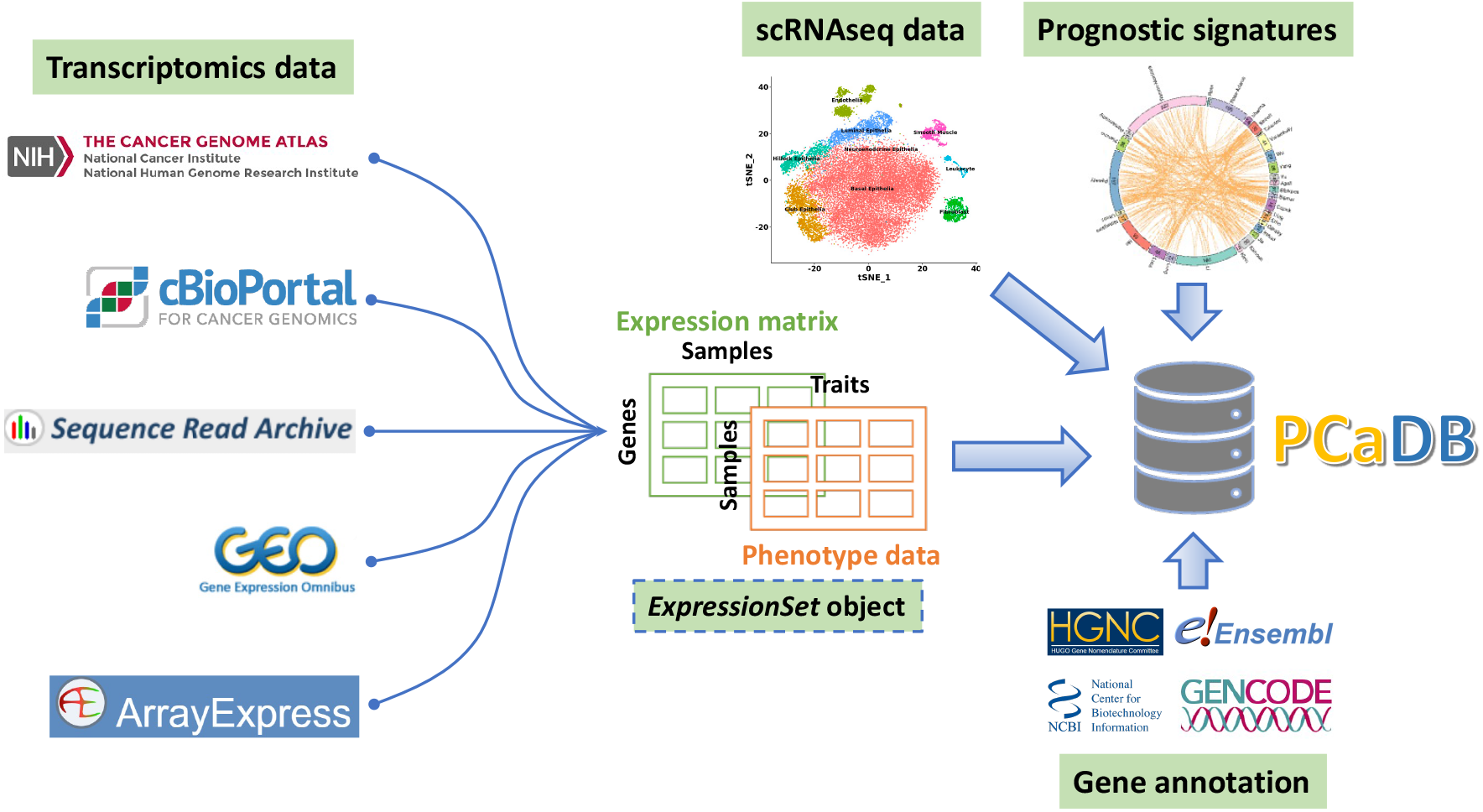
Overview of the collection of bulk transcriptomics data, scRNAseq data, published gene expression prognostic signatures, and gene annotation data for PCaDB.

## Materials and Methods

To obtain a complete list of transcriptomics data of PCa patient cohort, we conducted a comprehensive search in public data repositories, including the National Cancer Institute (NCI) Genomic Data Commons (GDC) (7), cBioportal (4), ArrayExpress (8), National Center for Biotechnology Information (NCBI) Gene Expression Omnibus (GEO) (9) and Sequence Read Archive (SRA) (10). The identification of datasets in GDC and cBioPortal were straightforward since the relevant data are grouped by cancer types. The keywords ‘prostate cancer’, ‘prostate tumor’, or ‘PCa’, AND ‘gene expression’, ‘mRNA’, ‘RNAseq’, ‘transcriptomics’, or ‘transcriptome’ were used to search the ArrayExpress, GEO, and SRA databases. Many other datasets were identified in previous publications (11–14). Datasets for cell lines or laboratory models were not included. Eventually, a total of 77 PCa transcriptomics datasets with 9,068 patient samples have been identified and deposited in PCaDB. Details about these datasets are summarized in Supplementary Table S1.

A single-cell RNAseq dataset for cell types isolated by fluorescence-activated cell sorting (FACS) from normal human prostates (5) was downloaded from the GenitoUrinary Development Molecular Anatomy Project (GUDMAP) database (15) and integrated into the PCaDB database, allowing for the investigation of gene expression at the single-cell level.

The gene lists of 30 published prognostic signatures for PCa, which were comprehensively evaluated in a previous study (6), were included in the PCaDB database for a more detailed functional characterization and prognostic performance comparison in each of the transcriptomics datasets.

We collected the gene annotation from the Ensembl (16), GENCODE (17), HUGO Gene Nomenclature Committee (HGNC) (18), and NCBI Entrez Gene (19) databases and developed a pipeline to map the different types of the gene identifiers including the Ensembl ID, HGNC approved gene symbol and alias symbol, as well as Entrez ID to facilitate the harmonization of transcriptomics data collected from different resources and the query of genes in PCaDB.

A suite of bioinformatics pipelines has been developed to download, process, and harmonize the gene expression and sample metadata from different resources. RNA sequencing data from SRA and Affymetrix microarray data from GEO with the raw FASTQ or CEL files were reprocessed using respective unified pipelines. For some other datasets with no raw data available, the normalized data were downloaded directly from the repositories. The methods for data processing described in the original publications were carefully inspected to make sure that appropriate processing methods had been used. Details about the data processing pipelines and database implementation can be found in the Supplementary Materials.

## Results

PCaDB provides a user-friendly interface and a suite of analytical and visualization tools for the comprehensive analysis of gene expression data at three levels: (*i*) query an individual gene of interest, (*ii*) characterization of prognostic signatures, and (*iii*) whole-transcriptome data analysis. Multiple analytical and visualization functions can be implemented in each module (Figure 2). The introduction to the functional modules is provided below. Details about the input data, bioinformatics tools, statistical algorithms, and parameter settings, etc. for each data analysis or visualization function can be found in the ‘Tutorial’ page in PCaDB.

**Figure 2.**
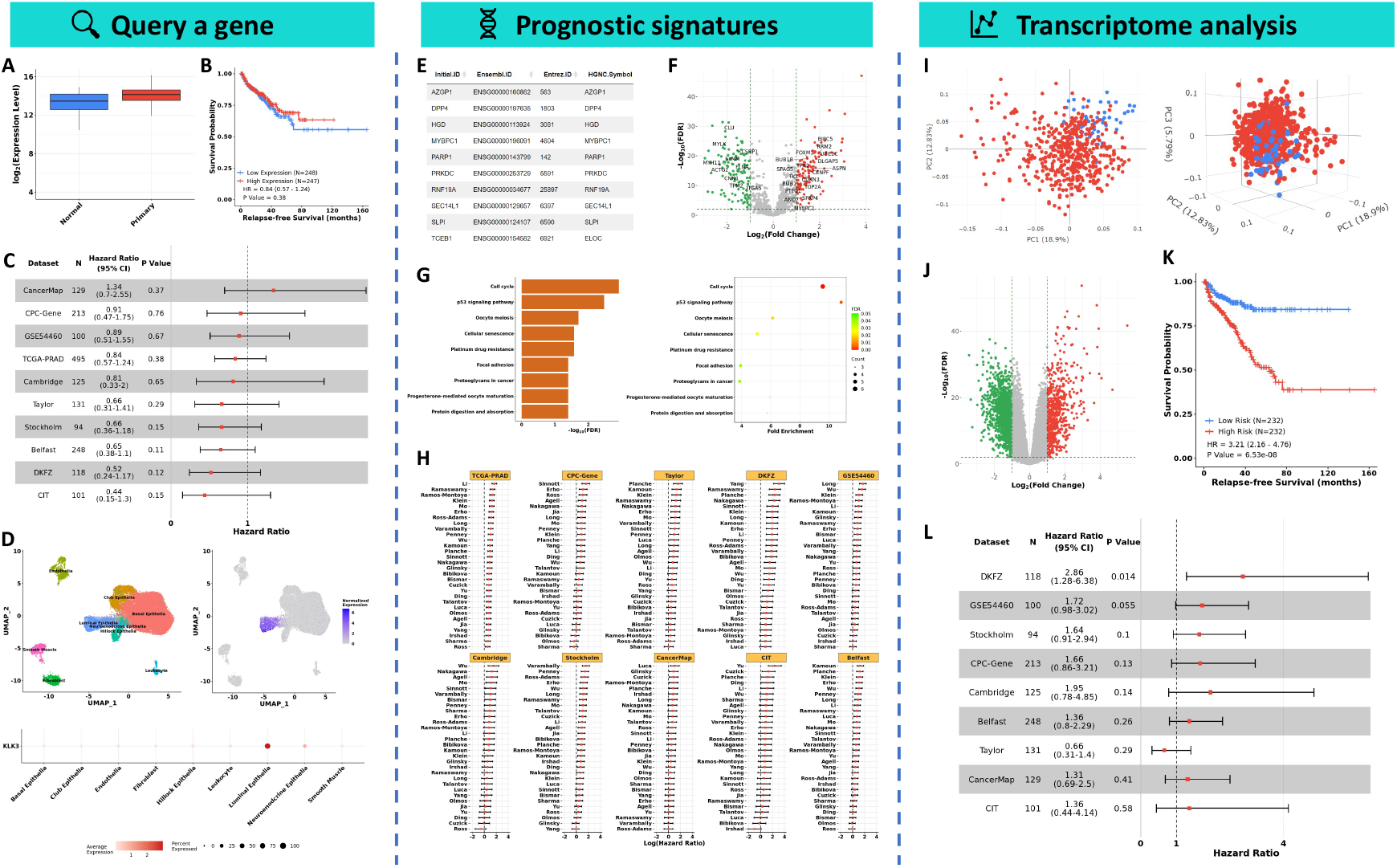
Comprehensive analysis of gene expression data at three levels in PCaDB, including the query of an individual gene of interest, characterization of prognostic signatures, and whole-transcriptome data analysis. (**A**) The box plot to visualize the expression of the gene of interest in different sample types. (**B**) The Kaplan Meier (KM) survival curve of relapse-free survival (RFS) for the selected gene in an individual dataset. (**C**) The forest plot to visualize the association of a gene of interest with RFS across multiple datasets. (**D**) Visualization of the gene expression in different prostate cell types using Uniform Manifold Approximation and Projection (UMAP) plot and bubble plot. (**E**) The list of genes in a selected prognostic signature. (**F**) Differential gene expression analysis of genes in all the prognostic signatures between tumor and normal samples. (**G**) Visualization of the functional enrichment analysis results using bar plot and bubble plot. (**H**) Comprehensive evaluation of the performances of all the prognostic signatures across multiple datasets based a selected training dataset and a selected machine learning algorithm. (**I**) 2D and 3D interactive visualization of the principal component analysis (PCA) result for a transcriptomics dataset. (**J**) Differential gene expression analysis for the whole transcriptome profiling data between case (e.g., primary tumor) and control (e.g., tumor-adjacent normal) samples. (**K**) Development of a prognostic model based on a user-provided gene list and a selected survival analysis algorithm. A KM survival curve is generated to evaluate the performance of the prognostic model in the training dataset. (**L**) Validation of the prognostic model leveraging gene expression data from multiple independent cohorts. The sample size, hazard ratio (HR), 95% confidence intervals (CIs), and p value for each validation cohort can be visualized in a forest plot.

### I. Query an individual gene of interest

Users can query a gene of interest by typing the Ensembl ID, NCBI Entrez ID, or HGNC approved symbol and alias symbol in the ‘Search a gene’ field and selecting the gene from the dropdown list. The general information about the gene and some useful external links to the databases, such as ENSEMBL, HGNC, and NCBI for more detailed description of the gene, Genotype-Tissue Expression (GTEx) (20) and Human Protein Atlas (HPA) (21) for the gene expression pattern in different human tissues, and Kyoto Encyclopedia of Genes and Genomes (KEGG) (22) for the pathways that the gene involves in are provided. A suite of advanced analyses and visualizations can be interactively performed for the selected gene, including differential expression (DE) analysis between different types of samples, RFS survival analysis across multiple cohorts, and gene expression analysis in different prostate cell types at the single-cell level.

The box plot is used to visualize the gene expression in different sample types, such as healthy, tumor-adjacent normal, primary tumor, or metastatic tumor in different tissues, depending on the availability of the data in the selected dataset (Figure 2A). Kaplan Meier (KM) analysis of relapse-free survival (RFS) can be performed in the 10 datasets with 1,558 primary tumor samples from PCa patients with the data of biochemical recurrence (BCR) status and follow up time after treatment (Figure 2B). A forest plot with the result from survival analysis, including the numbers of samples, hazard ratios (HRs), 95% confidence intervals (CIs), and p values across all the datasets, will be generated, and the KM survival curve for each dataset will also be plotted (Figure 2C). The expression pattern of the selected gene in different cell types from normal human prostate can be visualized using the pre-calculated t-distributed stochastic neighbor embedding (t-SNE) plot and uniform manifold approximation and projection (UMAP) plot, violin plot, and bubble plot with the average gene expression and percent of cells expressed for each cell type (Figure 2D).

### II. Characterization of prognostic signatures

A comprehensive evaluation of the prognostic performances of 30 published signatures was performed in a previous study and we included all those signatures in the PCaDB database, allowing for a more detailed characterization of the signatures, including DE analysis of the signature genes, KM survival analysis of RFS, functional enrichment analysis, and evaluation of the prognostic performances of the signatures across multiple cohorts.

On the ‘Prognostic Signatures’ page, the list of genes in a signature of interest can be viewed by selecting the signature from the dropdown list. The DE analysis of the signature genes between primary tumor and tumor-adjacent normal samples can be performed using the R package *limma* (23). A data table will be created to list the differentially expressed genes (DEGs) and a volcano plot will be generated to visualize these DEGs (Figures 2E and 2F). The KM survival analysis of RFS for the signature genes can be performed in each of the 10 datasets with RFS information. A forest plot for the common genes in 3 or more signatures and a data table with the survival analysis result for all the signature genes are generated. Functional enrichment analysis of the signature genes can be performed using the R package *clusterProfiler* (24). A data table is produced to summarize the significantly enriched pathways/ontologies. A bar plot and a bubble plot are used to visualize the top enriched pathways/ontologies based on the p values adjusted by the Benjamini-Hochberg (BH) method (Figure 2G). A more detailed evaluation of the performances of the signatures can be performed in PCaDB using different survival analysis algorithms and different training and test datasets. Users can select ‘All Signatures’ from the dropdown list to perform a comprehensive analysis to compare and rank the signatures in each test set based on three metrics, including concordance index (C-index), time-dependent receiver operating characteristics (ROC) curve, and HR estimated by the KM survival analysis (Figure 2H). If a given signature is selected, a prognostic model can be developed using the expression data of the signature genes in the selected training dataset and the selected survival analysis method. The risk score of each patient in the test datasets will be computed based on the model, and the C-indexes, the area under the ROC curves (AUCs), and the HRs are calculated to assess the prognostic power for the signature based on the independent test cohorts. Forest plots are used to visualize the results, while data tables with more detailed results are also provided.

### III. Whole-transcriptome data analysis

More advanced and comprehensive analyses can be performed at the whole-transcriptome level in PCaDB, allowing users to identify DEGs between different types of samples, discover biomarkers associated with clinical outcomes (*i*.*e*., BCR), as well as develop and validate gene expression-based signatures and models for PCa prognosis.

A transcriptome dataset of interest can be selected on the ‘Transcriptome Analysis’ page, and the summary of the dataset including platform, data processing pipeline, and the available metadata, such as sample type, preoperative prostate-specific antigen (PSA), Gleason score, BCR status, and time to BCR, will be displayed automatically. Principal component analysis (PCA) can be performed for dimensionality reduction to have a global view of the transcriptomics dataset, and a 2D or 3D interactive plot based on the first two or three principal components, respectively, will be generated for visualization (Figure 2I). The DE analysis using the whole-transcriptome data allows users to identify DEGs between many different types of samples by comparing the case and control groups. PCaDB has included a wide spectrum of sample types, which allows for answering many critical research questions at molecular level through the comparisons of samples of different types, such as localized or advanced tumor vs. tumor-adjacent normal samples, non-metastatic castration-resistant prostate cancer (CRPC) vs. metastatic CRPC (mCRPC), mCRPC at different sites (lymph node, bone, liver, lung, etc.), normal and tumor samples between African American men (AAM) vs. European American men (EAM), tumor samples collected pre-androgen-deprivation therapy (ADT) vs. those collected post-ADT, ADT responsive vs. castration resistant, etc. The R package *limma* (23) is used to identify DEGs in PCaDB (Figure 2J). Both the univariate Cox proportional hazards (CoxPH) and KM survival analyses of RFS can be performed at the whole-transcriptome level to identify biomarkers associated with clinical outcome of PCa in a selected dataset of interest. Multivariate CoxPH analysis for each individual gene can be also performed, with the important clinical factors, including the age at diagnosis, PSA level, Gleason score, and clinical T stage being adjusted. The biomarkers that are significant across multiple datasets may be used alone or in combination with other biomarkers to derive a prognostic signature for PCa. In PCaDB, users can provide a list of genes and select any survival analysis method, such as CoxPH, Cox model regularized with ridge penalty (Cox-Ridge), or lasso penalty (Cox-Lasso) (25), to develop a prognostic model using the selected dataset as a training set. Risk scores for the patients in the training set are calculated and the median value is used as the threshold to dichotomize these patients into low- and high-risk groups. A KM survival curve is generated to show the prognostic performance of the signature in the training set (Figure 2K). Similarly, risk scores are calculated for patients in each of the nine remaining datasets with RFS data, and a forest plot is generated to validate the prognostic model in independent cohorts (Figure 2L).

### Discussion and future directions

PCaDB is a comprehensive database for transcriptomes of PCa cohorts with a total of 9,068 samples from 77 public datasets. A suite of well-designed functions is provided in PCaDB for the interactive analysis and visualization of the transcriptomics data. A scRNAseq dataset for normal human prostates is also included, allowing for the investigation of gene expression at the single-cell level. Detailed characterization and evaluation of 30 published prognostic signatures can be performed in PCaDB to identify the most promising ones for further validations in prospective clinical studies. All the data have been processed with a comprehensive workflow, and the data can be easily downloaded from the database, allowing researchers with deep expertise in bioinformatics and data science to perform more advanced analyses to better understand the molecular basis of PCa initiation and progression, elucidate the mechanisms of therapeutic resistance, identify drug targets or potential drug combinations, and develop novel statistical methodologies or molecular signatures for PCa diagnosis, prognosis and treatment response prediction. All the pipelines developed in the study are publicly available, which can be used to promote the collection, harmonization, and analysis of gene expression profiling data for other cancer types. Comprehensive tutorials have been provided in the ‘PCaDB Pipeline’ and ‘Tutorial’ pages, allowing users to better understand the bioinformatics pipelines and analytical methods that are being used in PCaDB.

While PCaDB is diligently serving the prostate cancer research community, new datasets including other omics data and analytical methods will be included in PCaDB as soon as they are available. We expect that PCaDB would become a valuable online resource for a comprehensive analysis of PCa transcriptomics data to facilitate research studies and improve the clinical management of PCa.

## Availability of data and materials

The web interface to PCaDB is publicly available at http://bioinfo.jialab-ucr.org/PCaDB/. All the processed data can be downloaded on the ‘Download’ page of the database. The pipelines used for data downloading, processing, and harmonization are all available at https://github.com/rli012/PCaDB and on the ‘PCaDB Pipeline’ page in the PCaDB database.

## Authors’ contributions

R.L. and Z.J. designed the project and wrote the manuscript. R.L. collected the public transcriptomics datasets and built the database. J.Z. and W.Z. reviewed and revised the manuscript. All authors read and approved the final manuscript.

## Notes

**Financial Support** This work was supported by Z.J.’s UC Riverside Faculty Start-up Fund and UC Cancer Research Coordinating Committee Competition Award. J.Z. was supported by the Science and Technology Project of Guizhou Province in 2017 ([2017]5803), the High-level innovative talent project of Guizhou Province in 2018 ([2018]5639), and the Science and Technology Plan Project of Guiyang in 2019 ([2019]2-15). W.Z. was supported by the grants from National Natural Science Foundation of China (82072813, 8157142) and Guangzhou Municipal Science and Technology Project (201803040001).

**Competing interests** The authors declare no potential conflicts of interest

### Competing Interest Statement

The authors have declared no competing interest.

## References

1. Lovén J, Orlando DA, Sigova AA, Lin CY, Rahl PB, Burge CB, et al. Revisiting Global Gene Expression Analysis. Cell. 2012;151:476–82.

2. Sung H, Ferlay J, Siegel RL, Laversanne M, Soerjomataram I, Jemal A, et al. Global Cancer Statistics 2020: GLOBOCAN Estimates of Incidence and Mortality Worldwide for 36 Cancers in 185 Countries. CA: A Cancer Journal for Clinicians. 2021;71:209–49.

3. Tang Z, Li C, Kang B, Gao G, Li C, Zhang Z. GEPIA: a web server for cancer and normal gene expression profiling and interactive analyses. Nucleic Acids Research. 2017;45:W98–102.

4. Gao J, Aksoy BA, Dogrusoz U, Dresdner G, Gross B, Sumer SO, et al. Integrative analysis of complex cancer genomics and clinical profiles using the cBioPortal. Sci Signal. 2013;6:pl1.

5. Henry GH, Malewska A, Joseph DB, Malladi VS, Lee J, Torrealba J, et al. A Cellular Anatomy of the Normal Adult Human Prostate and Prostatic Urethra. Cell Reports. 2018;25:3530-3542.e5.

6. Li R, Zhu J, Zhong W-D, Jia Z. Comprehensive evaluation of machine learning models and gene expression signatures for prostate cancer prognosis using large population cohorts. bioRxiv. 2021. doi.org/10.1101/2021.07.02.450975.

7. Jensen MA, Ferretti V, Grossman RL, Staudt LM. The NCI Genomic Data Commons as an engine for precision medicine. Blood. 2017;130:453–9.

8. Sarkans U, Füllgrabe A, Ali A, Athar A, Behrangi E, Diaz N, et al. From ArrayExpress to BioStudies. Nucleic Acids Research. 2021;49:D1502–6.

9. Edgar R, Domrachev M, Lash AE. Gene Expression Omnibus: NCBI gene expression and hybridization array data repository. Nucleic Acids Research. 2002;30:207–10.

10. Katz K, Shutov O, Lapoint R, Kimelman M, Brister JR, O’Sullivan C. The Sequence Read Archive: a decade more of explosive growth. Nucleic Acids Research. 2021;gkab1053.

11. Li R, Wang S, Cui Y, Qu H, Chater JM, Zhang L, et al. Extended application of genomic selection to screen multiomics data for prognostic signatures of prostate cancer. Brief Bioinform. 2021;22:bbaa197.

12. Yang L, Roberts D, Takhar M, Erho N, Bibby BAS, Thiruthaneeswaran N, et al. Development and Validation of a 28-gene Hypoxia-related Prognostic Signature for Localized Prostate Cancer. EBioMedicine. 2018;31:182–9.

13. You S, Knudsen BS, Erho N, Alshalalfa M, Takhar M, Al-Deen Ashab H, et al. Integrated Classification of Prostate Cancer Reveals a Novel Luminal Subtype with Poor Outcome. Cancer Res. 2016;76:4948–58.

14. Luca B-A, Moulton V, Ellis C, Connell SP, Brewer DS, Cooper CS. Convergence of Prognostic Gene Signatures Suggests Underlying Mechanisms of Human Prostate Cancer Progression. Genes (Basel). 2020;11:E802.

15. Harding SD, Armit C, Armstrong J, Brennan J, Cheng Y, Haggarty B, et al. The GUDMAP database--an online resource for genitourinary research. Development. 2011;138:2845–53.

16. Howe KL, Achuthan P, Allen J, Allen J, Alvarez-Jarreta J, Amode MR, et al. Ensembl 2021. Nucleic Acids Research. 2021;49:D884–91.

17. Frankish A, Diekhans M, Jungreis I, Lagarde J, Loveland JE, Mudge JM, et al. GENCODE 2021. Nucleic Acids Research. 2021;49:D916–23.

18. Tweedie S, Braschi B, Gray K, Jones TEM, Seal RL, Yates B, et al. Genenames.org: the HGNC and VGNC resources in 2021. Nucleic Acids Research. 2021;49:D939–46.

19. Maglott D, Ostell J, Pruitt KD, Tatusova T. Entrez Gene: gene-centered information at NCBI. Nucleic Acids Research. 2005;33:D54–8.

20. The GTEx Consortium TGte. The GTEx Consortium atlas of genetic regulatory effects across human tissues. Science. American Association for the Advancement of Science; 2020;369:1318–30.

21. Uhlen M, Oksvold P, Fagerberg L, Lundberg E, Jonasson K, Forsberg M, et al. Towards a knowledge-based Human Protein Atlas. Nat Biotechnol. 2010;28:1248–50.

22. Kanehisa M, Furumichi M, Tanabe M, Sato Y, Morishima K. KEGG: new perspectives on genomes, pathways, diseases and drugs. Nucleic Acids Research. 2017;45:D353–61.

23. Ritchie ME, Phipson B, Wu D, Hu Y, Law CW, Shi W, et al. limma powers differential expression analyses for RNA-sequencing and microarray studies. Nucleic Acids Research. 2015;43:e47–e47.

24. Wu T, Hu E, Xu S, Chen M, Guo P, Dai Z, et al. clusterProfiler 4.0: A universal enrichment tool for interpreting omics data. The Innovation. 2021;100141.

25. Simon N, Friedman J, Hastie T, Tibshirani R. Regularization Paths for Cox’s Proportional Hazards Model via Coordinate Descent. J Stat Softw. 2011;39:1– 13.

