## Supplementary Materials for "PCaDB - a comprehensive and interactive database for transcriptomes from prostate cancer population cohorts"

### Supplementary Materials and Methods

#### Data processing for the bulk transcriptomics data, sample metadata, and single-cell RNAseq data from public data repositories

A comprehensive workflow has been developed to download, process, and harmonize the transcriptomics data and the associated sample metadata from public data repositories, including National Cancer Institute (NCI) Genomic Data Commons (GDC) for The Cancer Genome Atlas Prostate Adenocarcinoma (TCGA-PRAD) data (1,2), cBioportal (3), ArrayExpress (4), National Center for Biotechnology Information (NCBI) Gene Expression Omnibus (GEO) (5) and Sequence Read Archive (SRA) (6). In general, gene expression profiling data generated by RNA sequencing (RNAseq) and Affymetrix microarray platforms were reprocessed if raw data (*i.e.*, FASTQ or CEL files) were available. Otherwise, the normalized data such as the fragments/reads per kilobase of transcript per million mapped reads (FPKM/RPKM) values for RNAseq data and normalized intensities for microarray data were downloaded directly from the corresponding data repositories. For the small proportion of datasets that only have normalized data provided in the public data repositories, the methods for data processing described in the original papers were carefully inspected to make sure that appropriate processing methods had been used. All the gene expression data in PCaDB are in log2 scale and the Ensembl gene identifiers are used for all the datasets. If multiple probes/genes matched to the same Ensembl ID, only the most informative one with the maximum interquartile range (IQR) for the gene expression was used for this Ensembl ID. Details about the pipelines for downloading, processing, and harmonization of gene expression data generated using different platforms or obtained from different repositories are described below.

(i) RNAseq data from SRA

The *fasterq-dump* tool in SRA Toolkit (2.10.8) (https://github.com/ncbi/sra-tools) was used to download and convert SRA data into FASTQ format for the RNAseq data in SRA. *FastQC* (v0.11.5) (https://www.bioinformatics.babraham.ac.uk/projects/fastqc/) was used for a comprehensive quality control (QC) of the sequencing data, including base sequence quality, GC content, overrepresented sequences, and adapter content, etc. *STAR* (2.7.9a) (7) was adopted to align the sequencing reads to the reference genome GRCh38 (primary assembly) based on the GENCODE (release 38) gene annotation (8). The 2-pass mapping was performed for each sample separately. The BAM files generated from the alignment step were sorted using *samtools* (v1.9) (9). The *featureCounts* program in the *Subread* package (10) was used for gene expression quantification to generate the count matrix. Only read pairs that were uniquely mapped to the exonic regions were quantified. The count data was normalized using the Trimmed Mean of M values (TMM) method implemented in the R package *edgeR* (11), and lowly-expressed genes with counts per million (CPM) < 1 in more than 50% of samples in the dataset were filtered out prior to any downstream analyses. To generate a more comprehensive QC report, we used the *rnaseqc* (v2.4.2) tool (12) to characterize the quality of the RNA, sequencing data, alignment, and expression profile of each sample. Finally, an aggregated QC report for each dataset was generated using *MultiQC* (13) to display the QC metrics from multiple bioinformatics analyses, including *FastQC*, *STAR*, *featureCounts*, and *rnaseqc*.

(ii) RNAseq data from TCGA-PRAD

The HTSeq-Counts data for TCGA-PRAD project is publicly available in GDC, which was downloaded and processed by a series of functions in the R package *GDCRNATools* (14) to generate the count matrix. The count data was also normalized using the TMM method implemented in the R package *edgeR* (11). Similarly, lowly-expressed genes with CPM < 1 in more than 50% of samples in the dataset were filtered out prior to any downstream analyses.

(iii) RNAseq and microarray data from cBioPortal

Although transcriptomics data for multiple PCa cohorts have been included in cBioPortal, the cBioPortal data were usually not used in PCaDB, unless the raw data were not available in any of the other data repositories, because only the normalized data were provided in cBioPortal, which may not be suitable for some of the downstream analyses. e.g., the FPKM/RPKM values are not recommended for DE analysis. For the few datasets from cBioPortal, normalized gene expression data (FPKM/RPKM values for RNAseq data or normalized intensities for microarray data) were downloaded manually from cBioPortal. Log2 transformation may be performed if it hadn't been done and the gene symbols/entrez ids were all converted to Ensembl IDs. If multiple genes matched to the same Ensembl ID, only the most informative one with the maximum interquartile range (IQR) for the gene expression is used for this Ensembl ID.

(iv) Microarray data from GEO and ArrayExpress

Affymetrix arrays (e.g., Affymetrix Human Exon 1.0 ST Array, Affymetrix Human Gene 2.0 ST Array, Affymetrix Human Genome U133A Array, and Affymetrix Human Genome U133 Plus 2.0 Array, etc.) with raw CEL files available in GEO or ArrayExpress were reprocessed by the unified PCaDB pipeline. The R package *GEOquery* (15) was used to download the CEL files and the Robust Multichip Average (RMA) method implemented in the R package *oligo* (16) was used for background correction, quantile normalization, and log2 transformation. The annotation package downloaded from the Brainarray database (Version 24.0.0; GENCODE release 32) was used for probe/gene annotation (17). Raw CEL files in ArrayExpress were downloaded using a custom script and the expression data were reprocessed using the same pipeline as that for GEO data.

The only Affymetrix microarray dataset that was not reprocessed by our PCaDB pipeline is the CPC-Gene dataset (GSE107299). The reason was that the data was generated based on two different Affymetrix arrays: Affymetrix Human Gene 2.0 ST Array (Batch 1,3,4,5) and Affymetrix Human Transcriptome Array 2.0 (Batch 2) with a batch effect. Therefore, the normalized and batch-corrected data was downloaded directly from GEO. Likewise, the normalized intensities for the microarray datasets generated using other platforms (e.g., Illumina BeadArray or Agilent Whole Human Genome Oligo Microarray) were downloaded directly from the repository using *GEOquery*. The methods for data processing described in the original papers were carefully inspected to make sure that appropriate processing methods had been used. Log2 transformation may be performed on the normalized data if it hadn't been done and the probe/gene ids were all converted to Ensembl IDs. If multiple probes/genes matched to the same Ensembl ID, only the most informative one with the maximum IQR for the gene expression is used for this Ensembl ID.

(v) Sample metadata

Metadata associated with the samples were obtained from the public data repositories and harmonized using a custom script followed by a careful manual curation. The pipelines for metadata harmonization are very similar for data from different repositories. The main difference is the way to download and retrieve the data.

For sample metadata from TCGA-PRAD project, the XML files with the clinical information of the patients were downloaded and organized using the R package *GDCRNATools*. Some clinical features that are not available in the GDC data portal, such as preoperative prostate-specific antigen (PSA) level, were retrieved from Broad GDAC Firehose (https://gdac. broadinstitute.org/).

The phenotype data of datasets in GEO were retrieved using our in-house web tool - *WebGEO* (http://bioinfo.jialab-ucr.org/WebGEO/; under development) based on the R package *GEOquery*, which allows for querying and downloading the sample metadata of a dataset in seconds. In case users are interested in downloading phenotype data programmatically using *GEOquery*, the script is also provided in the 'PCaDB Pipeline' page in PCaDB. Sample metadata (.sdrf.txt files) for the ArrayExpress datasets were downloaded using a custom script.

Most of the SRA datasets also have corresponding GEO accessions, thus the sample metadata for these datasets were also retrieved from GEO using *WebGEO*. For a few SRA datasets that do not have corresponding GEO accessions, the metadata (SraRunTable) were downloaded manually from SRA. Likewise, clinical data for the cBioPortal datasets were manually downloaded from the portal.

In addition, the original publications were carefully reviewed to further ensure the accuracy and comprehensiveness of the sample metadata for each dataset which has been included in PCaDB.

We have created a comprehensive list of 33 field names, including sample id, patient id, tissue, batch, sample type, age at diagnosis, ethnicity, race, clinical stages, pathology stages, preoperative PSA, Gleason score, overall survival, relapse-free survival, treatment, etc., for the sample metadata. The complete list of the field names can be found in the 'PCaDB Pipeline' page of PCaDB.

(vi) Single-cell RNAseq data for normal human prostates

For the scRNAseq dataset, the *Seurat* object can be downloaded directly from the GUDMAP (https://www.gudmap.org/chaise/record/#2/RNASeq:Study/RID=W-RAHW) (18). The normalized gene expression matrix, cell type annotation, as well as the t-distributed stochastic neighbor embedding (t-SNE) and uniform manifold approximation and projection (UMAP) coordinates were included in an R list object. Details about the scRNAseq, such as data processing, data normalization, t-SNE and UMAP analyses, etc., have been described in the original study (19).

**Integration of gene annotation data**

It's very critical to map different types of gene identifiers from different databases for the data analysis in a transcriptome atlas and for the integrated analysis of multiple datasets, especially when the data were generated using different platforms, processed with different bioinformatics pipelines, or collected from different resources. The Ensembl gene annotation (Release 105) was downloaded using the R package *biomaRt* (20). The GTF file for the GENCODE gene annotation (Release 38) was downloaded from the FTP site (http://ftp.ebi.ac.uk/pub/databases/gencode/Gencode_human/release_38/). The two files hgnc_complete_set.txt and withdrawn.txt for the HGNC gene annotation were downloaded from the FTP site (http://ftp.ebi.ac.uk/pub/databases/genenames/new/tsv/), and the two files gene_info.gz and gene_history.gz for the NCBI gene annotation were downloaded from the FTP site (https://ftp.ncbi.nih.gov/gene/DATA/). A comprehensive pipeline was developed to map the gene IDs and the source code is publicly available at https://github.com/rli012/PCaDB and on the 'PCaDB Pipeline' page in the database.

### Database implementation

PCaDB has been developed using R Shiny (https://CRAN.R-project.org/package=shiny), which provides an elegant and powerful web framework for building interactive web applications using the R language (https://www.R-project.org/). The major advantage of Shiny is that a lot of R/Bioconductor packages such as *limma* (21), *clusterProfiler* (22), *Biobase* (23), *ggplot2* (24), etc. can be used for the advanced bioinformatics analyses and visualization. In PCaDB, the majority of the visualizations are based on the R package *ggplot2*, interactive tables are generated using the R package *DT* (https://CRAN.R-project.org/package=DT), allowing users to filter, sort, copy, and download the data, and interactive plots are made using the R package *plotly* (25). The blue gradient theme in the *dashboardthemes* package (https://CRAN.R-project.org/package=dashboardthemes) is used with some modifications. The web application is deployed on Amazon Web Services (AWS).

### Data download module in PCaDB

All the harmonized data, including the 77 public PCa transcriptomics datasets, the scRNAseq data for normal human prostates, the summary and the gene lists of the 30 published prognostic signatures, and the integrated gene annotation data, can be downloaded easily on the 'Download' page of PCaDB. The summary of the transcriptomics datasets including the sample size, the GEO, ArrayExpress, or SRA accession number, the gene expression profiling platform, the bioinformatics pipeline that was used to process the data, etc. is also available for downloading. The *ExpressionSet* class is used for the gene expression data and sample metadata of the transcriptomics datasets, and the data can be downloaded in the RDS format. The *Seurat* object of the scRNAseq data and the gene annotation data are also available for users to download in the RDS format.

### References


1. Jensen MA, Ferretti V, Grossman RL, Staudt LM. The NCI Genomic Data Commons as an engine for precision medicine. Blood. 2017;130:453–9.
2. Abeshouse A, Ahn J, Akbani R, Ally A, Amin S, Andry CD, et al. The Molecular Taxonomy of Primary Prostate Cancer. Cell. 2015;163:1011–25.
3. Gao J, Aksoy BA, Dogrusoz U, Dresdner G, Gross B, Sumer SO, et al. Integrative analysis of complex cancer genomics and clinical profiles using the cBioPortal. Sci Signal. 2013;6:pl1.
4. Sarkans U, Füllgrabe A, Ali A, Athar A, Behrangi E, Diaz N, et al. From ArrayExpress to BioStudies. Nucleic Acids Research. 2021;49:D1502–6.
5. Barrett T, Wilhite SE, Ledoux P, Evangelista C, Kim IF, Tomashevsky M, et al. NCBI GEO: archive for functional genomics data sets—update. Nucleic Acids Research. 2013;41:D991–5.
6. Katz K, Shutov O, Lapoint R, Kimelman M, Brister JR, O'Sullivan C. The Sequence Read Archive: a decade more of explosive growth. Nucleic Acids Research. 2021;gkab1053.
7. Dobin A, Davis CA, Schlesinger F, Drenkow J, Zaleski C, Jha S, et al. STAR: ultrafast universal RNA-seq aligner. Bioinformatics. 2013;29:15–21.
8. Frankish A, Diekhans M, Jungreis I, Lagarde J, Loveland JE, Mudge JM, et al. GENCODE 2021. Nucleic Acids Research. 2021;49:D916–23.
9. Li H, Handsaker B, Wysoker A, Fennell T, Ruan J, Homer N, et al. The Sequence Alignment/Map format and SAMtools. Bioinformatics. 2009;25:2078–9.
10. Liao Y, Smyth GK, Shi W. featureCounts: an efficient general purpose program for assigning sequence reads to genomic features. Bioinformatics. 2014;30:923–30.
11. Robinson MD, McCarthy DJ, Smyth GK. edgeR: a Bioconductor package for differential expression analysis of digital gene expression data. Bioinformatics. 2010;26:139–40.
12. Graubert A, Aguet F, Ravi A, Ardlie KG, Getz G. RNA-SeQC 2: efficient RNA-seq quality control and quantification for large cohorts. Bioinformatics. 2021;37:3048–50.
13. Ewels P, Magnusson M, Lundin S, Käller M. MultiQC: summarize analysis results for multiple tools and samples in a single report. Bioinformatics. 2016;32:3047–8.

14. Li R, Qu H, Wang S, Wei J, Zhang L, Ma R, et al. GDCRNATools: an R/Bioconductor package for integrative analysis of lncRNA, miRNA and mRNA data in GDC. Bioinformatics. 2018;34:2515–7.
15. Davis S, Meltzer PS. GEOquery: a bridge between the Gene Expression Omnibus (GEO) and BioConductor. Bioinformatics. 2007;23:1846–7.
16. Carvalho BS, Irizarry RA. A framework for oligonucleotide microarray preprocessing. Bioinformatics. 2010;26:2363–7.
17. Dai M, Wang P, Boyd AD, Kostov G, Athey B, Jones EG, et al. Evolving gene/transcript definitions significantly alter the interpretation of GeneChip data. Nucleic Acids Research. 2005;33:e175–e175.
18. Harding SD, Armit C, Armstrong J, Brennan J, Cheng Y, Haggarty B, et al. The GUDMAP database--an online resource for genitourinary research. Development. 2011;138:2845–53.
19. Henry GH, Malewska A, Joseph DB, Malladi VS, Lee J, Torrealba J, et al. A Cellular Anatomy of the Normal Adult Human Prostate and Prostatic Urethra. Cell Reports. 2018;25:3530-3542.e5.
20. Durinck S, Spellman PT, Birney E, Huber W. Mapping identifiers for the integration of genomic datasets with the R/Bioconductor package biomaRt. Nat Protoc. 2009;4:1184–91.
21. Ritchie ME, Phipson B, Wu D, Hu Y, Law CW, Shi W, et al. limma powers differential expression analyses for RNA-sequencing and microarray studies. Nucleic Acids Research. 2015;43:e47–e47.
22. Wu T, Hu E, Xu S, Chen M, Guo P, Dai Z, et al. clusterProfiler 4.0: A universal enrichment tool for interpreting omics data. The Innovation. 2021;100141.
23. Huber W, Carey VJ, Gentleman R, Anders S, Carlson M, Carvalho BS, et al. Orchestrating high-throughput genomic analysis with Bioconductor. Nat Methods. 2015;12:115–21.
24. Wickham H. ggplot2: Elegant Graphics for Data Analysis. New York: Springer-Verlag; 2009. https://www.springer.com/gp/book/9780387981413
25. Sievert C. Interactive Web-Based Data Visualization with R, plotly, and shiny. CRC Press; 2020.
